# Nerves innervating copulatory organs show common FMRFamide, FVRIamide, MIP and serotonin immunoreactivity patterns across Dinophilidae (Annelida) indicating their conserved role in copulatory behaviour

**DOI:** 10.1101/491886

**Authors:** Alexandra Kerbl, Emilie Winther Tolstrup, Katrine Worsaae

## Abstract

**Background:** Males of the microscopic annelid family Dinophilidae use their prominent glandomuscular copulatory organ (penis) to enzymatically dissolve the female’s epidermis and thereafter inject sperm. In order to test for putative conserved copulatory structures and neural orchestration across three dinophilid species, we reconstructed the reproductive myo- and neuroanatomy and mapped neurotransmitter immunoreactivity patterns of two specific markers with reported roles in invertebrate male mating behaviour (FVRIamide, MIP) and three general neural markers (acetylated α-tubulin, serotonin, FMRFamide).

**Results:** Seminal vesicles (one or two pairs), surrounded by a thin layer of longitudinal and circular muscles and innervated by nerve fibres, are found between testes and copulatory organ in the larger males of *Dinophilus vorticoides* and *Trilobodrilus axi*, but are missing in the only 0.05 mm long *D. gyrociliatus* dwarf males. The midventral copulatory organ is in all species composed of an outer muscular penis sheath and an inner penis cone. Nerves encircle the organ equatorially, either as a ring-shaped circumpenial fibre mass or as dorsal and ventral commissures, which are connected to the ventrolateral nerve cords. All three examined dinophilids show similar serotonin-, FMRFamide-, and FVRIamide-like immunoreactivity patterns in the nerves surrounding the penis, supporting the general involvement of these neurotransmitters in copulatory behaviour in meiofaunal annelids. MIP-like immunoreactivity is restricted to the circumpenial fibre mass in *D. gyrociliatus* and commissures around the penis in *T. axi* (but not found in *D. vorticoides*), indicating its role in controlling the copulatory organ.

**Conclusions:** The overall myo- and neuroanatomy of the reproductive organs is rather similar in the three studied species, suggesting a common ancestry of the unpaired glandomuscular copulatory organ and its innervation in Dinophilidae. This is furthermore supported by the similar immunoreactivity patterns of the tested neurotransmitters around the penis. Smaller differences in the immunoreactivity patterns around the seminal vesicles and spermioducts might account for additional, individual traits. We thus show morphological support for the putatively conserved role of FMRFamide, FVRIamide, MIP and serotonin in dinophilid copulatory behaviour.

## Background

Invertebrates, including microscopic annelids living among sand grains (interstitially), show various forms of sexual reproduction, including hermaphroditism, and asexual multiplication such as budding or fissure [1,2]. Many interstitial groups, and among them the family Dinophilidae Macalister, 1876, show internal fertilization by direct sperm transfer, which diminishes loss of gametes and increases chances of successful reproduction [2–5]. Dinophilidae accomplish this by hypodermal injection, where the males enzymatically dissolve the epidermis of the females and transfer their sperm into the female’s body by pumping, both accomplished by their glandomuscular copulatory organ [4–8].

Dinophilidae comprises 18 strictly marine species, all characterized by small size, six segments, a dense and broad ventral ciliary band, a muscular pharynx, and complete lack of parapodia, chaetae, or other appendages [2,9]. The respective species can be distinguished by the number, continuity and arrangement of thin transverse ciliary bands, colour, life strategy as well as the basic architecture of the nervous system [10,11]. The dorsal brain does not show any compartmentalization such as lobes or ganglia, and the ventral nervous system is pentaneural with one pair of ventrolateral nerve cords, one medioventral nerve and in between these structures one or two pairs of paramedian nerves, which extend along the body. Furthermore, six main, ganglionated commissures connecting the ventrolateral cords signify the body segments, accompanied by one to several thin anterior commissures [10,11]. Despite their simple morphology, male dinophilids exhibit a relatively complex copulatory organ and show mating behaviour, which in the case of *Trilobodrilus axi* even includes a “mating dance” prior to copulation, where males repeatedly move their posterior body to the side when meeting females, or have been shown to crawl alongside each other [7].

Three different life strategies and major morphotypes have been described, exemplified by 1) the monomorphic and hyaline *Trilobodrilus axi* Westheide, 1967 (being 0.8-1.0mm long, and having a life cycle of approximately one year including a reproductive period between April and June [7,12,13]), 2) the monomorphic and yellow-orange pigmented *Dinophilus vorticoides* O. Schmidt, 1848 (having a body length of 1.5-2.5mm, a long life cycle including an extended, several months long encystment stage [6,14,15]), and 3) the sexually dimorphic and hyaline *Dinophilus gyrociliatus* O. Schmidt, 1857 (forming a diminutive 0.05mm long dwarf male in contrast to the 0.7-1.3mm long females, and having a short life cycle of three to four weeks [16–18]). Most dinophilids (with the exception of *T. hermaphroditus* Riser, 1999) are gonochoristic, with males having a muscular copulatory organ and associated penial glands [9]. Secretions produced by the latter enzymatically create a hole in the epidermis of the female, and sperm is then transferred into the female’s body cavity by one or more males [5,7,8,14,19,20]. Some *Dinophilus* species have stylet glands, which are described to aid the penetration of the female epidermis by keeping the wound open [5,8]. When fertilized, the eggs are released by momentary rupture of the epidermis. After release, the eggs become encapsulated in a cocoon-like gelatinous structure, which is either surrounding individual eggs or entire clutches. The animals develop directly, whereby hatching juveniles largely resemble the adults [2,9,12].

The male copulatory organ (penis) has been studied to some extend in members of the genus *Dinophilus* [4–8,14,21]. Especially in the dwarf male of *D. gyrociliatus*, where it takes up approximately one third of their entire body (with the testes filling most of the rest), the details of muscles and glands of the copulatory organ as well as the nerves possibly innervating it have been investigated by ultrastructural and immunohistochemical techniques [4,18,21], and especially Traut [8] conducted detailed behavioural observations. While there also have been histological observations of the reproductive organs in *D.* cf. *taeniatus* [5,6] (which now should be regarded as *D. vorticoides* based on its reported location on the Swedish coast (Gonzalez et al., in prep.)), *T. axi* and *T. heideri* [4,7], we lack a common three-dimensional morphological base in order to compare these organs more reliably. Despite the differences in size of the copulatory organ and the number of seminal vesicles between the species, the organization of the penis into a sheath and a cone or internal layer seems to be highly similar in all so far investigated species [4,5,18]. Comprehensive (immunohistochemical) studies have not been conducted on any of these species yet and the homology of the respective structures has not been addressed, although previous studies already hypothesized about functional similarities between at least the two *Dinophilus* species [4,8].

The nervous system and its connections to muscles and glands of the reproductive organs have hitherto only been studied in high detail in *D. gyrociliatus* [21,22]. Here, a pair of ganglia (consisting of four neurons each) is associated with the copulatory organ, and supposedly aids the integration of sensory cues from sensory cells of the posterior end of the animal (surrounding the gonopore) with the information from the anterior part of the body, as well as orchestrates the musculature and glands of the copulatory organ [18,21,22]. In contrast to these ganglia, the larger males of *D. vorticoides* and *T. axi* have not been reported to form specific penis ganglia in close proximity to the penis, but instead seem to show a more direct connection to the ventral nervous system [4,5]. However, details of the neural innervation of the copulatory organ as well as the presence and pattern of specific neurotransmitters in this local copulatory circuitry have not been studied previously.

Whereas pan-neural markers can be used to describe the nearly complete layout of the nervous system, neurotransmitters, which are only present in parts of it, can add more detailed information about putative functions of specific nervous system regions. Similar to other signalling molecules, they control behaviour by propagating, inhibiting, decreasing, or increasing action potentials of neurons, and thereby lead to excitation or inhibition of the innervated tissue [23]. Neurotransmitters can furthermore be classified into different categories based for example on their structure and formation, e.g., neuropeptides being chains of amino acids, and monoamines being downstream products of aromatic amino acids. Genetically conserved propeptide sequences coding for specific neuropeptides have been found to be correlated to orthologues across larger taxonomic groups (families, phyla) or even dating back to the origin of Bilateria [24–26]. Conzelmann *et al.* [24] demonstrated a conservation of the sequences as well as of the related immunoreactivities in specific cell types across Spiralia, thereby emphasizing the potential of antibodies against these neuropeptides to extrapolate functional subregions of nervous systems across animal groups. Several of these neuropeptides such as FMRFamide, FVRIamide and MIP (myoinhibitory peptide) have also been shown in functional studies (targeting especially molluscan species as well as ecdysozoans such as the model animal *Caenorhabditis elegans* [27–31]) to directly affect the musculature of the reproductive system. Despite this conservation, a previous comparative study of immunoreactivity patterns of 14 antibodies against specific propeptides found vast differences in the nervous systems and especially the brains of *D. gyrociliatus*, *D.*cf. *taeniatus* (which was collected in the Faroe Islands and therefore also should be referred to as *D. vorticoides* from now on, Gonzelez et al., in prep.) and *T. axi* [11]. The brains of all these species, however, contain several hundred cells, whose processes are tightly connected in the brain neuropil and are therefore potentially more difficult to analyse than the less compact local circuits underlying e.g. mating behaviour and reproduction. Focussing on the involvement of the tested neurotransmitters in the demarcated copulatory circuits of three closely related species will allow a better insight into the conservation of function across these closely related species, which will further aid to our understanding of neuroregulation in Spiralia.

In order to provide a detailed description of the neuromuscular anatomy of the male reproductive organs of the three dinophilid species *T. axi*, *D. vorticoides*, and *D. gyrociliatus*, we labelled musculature with phalloidin and nerves with antibodies against acetylated α-tubulin, the monoamine neurotransmitter serotonin, and three neuropeptides (FMRFamide, FVRIamide and MIP). Especially the latter have been described to play a role in mating/copulatory behaviour in invertebrates [27–31], mainly by orchestrating muscular contractions, amongst other functions they fullfill in the organisms. Whereas we used commercially available antibodies against acetylated α-tubulin, serotonin and FMRFamide, the antibodies against FVRIamide and MIP were customized against peptide sequences of *Platynereis dumerilii* (and show sufficient overlap with sequences of at least one of the tested species, as was shown by Kerbl *et al.* [11]) and *D. gyrociliatus* (see Materials and Methods for more details). This approach allowed us to compare the reproductive systems in great detail and discuss the evolution and possible functionality of their individual neural components. These hypotheses will be the base for future studies including behavioural assays, and receptor localization.

## Results

### Overview of the male reproductive system

In general, the reproductive systems in all three investigated dinophilid species consist of a medioventral glandomuscular copulatory organ or penis (co, Figs 1, 2, 3, 4) connected to the testes through spermioducts and one (in *Dinophilus vorticoides*) or two pairs of seminal vesicles (in *Trilobodrilus axi*, sv, Figs 1a-d, 2a, b, e, f, g). The latter are reduced to simple spermioducts in the dwarf males of *Dinophilus gyrociliatus*, and although their anterior portions resemble seminal vesicles due to their dense filling with sperm upon maturation, they lack the muscular lining and sphincter muscles observed in the other two species (Figs 3a, b, 4c).

**Figure 1.**
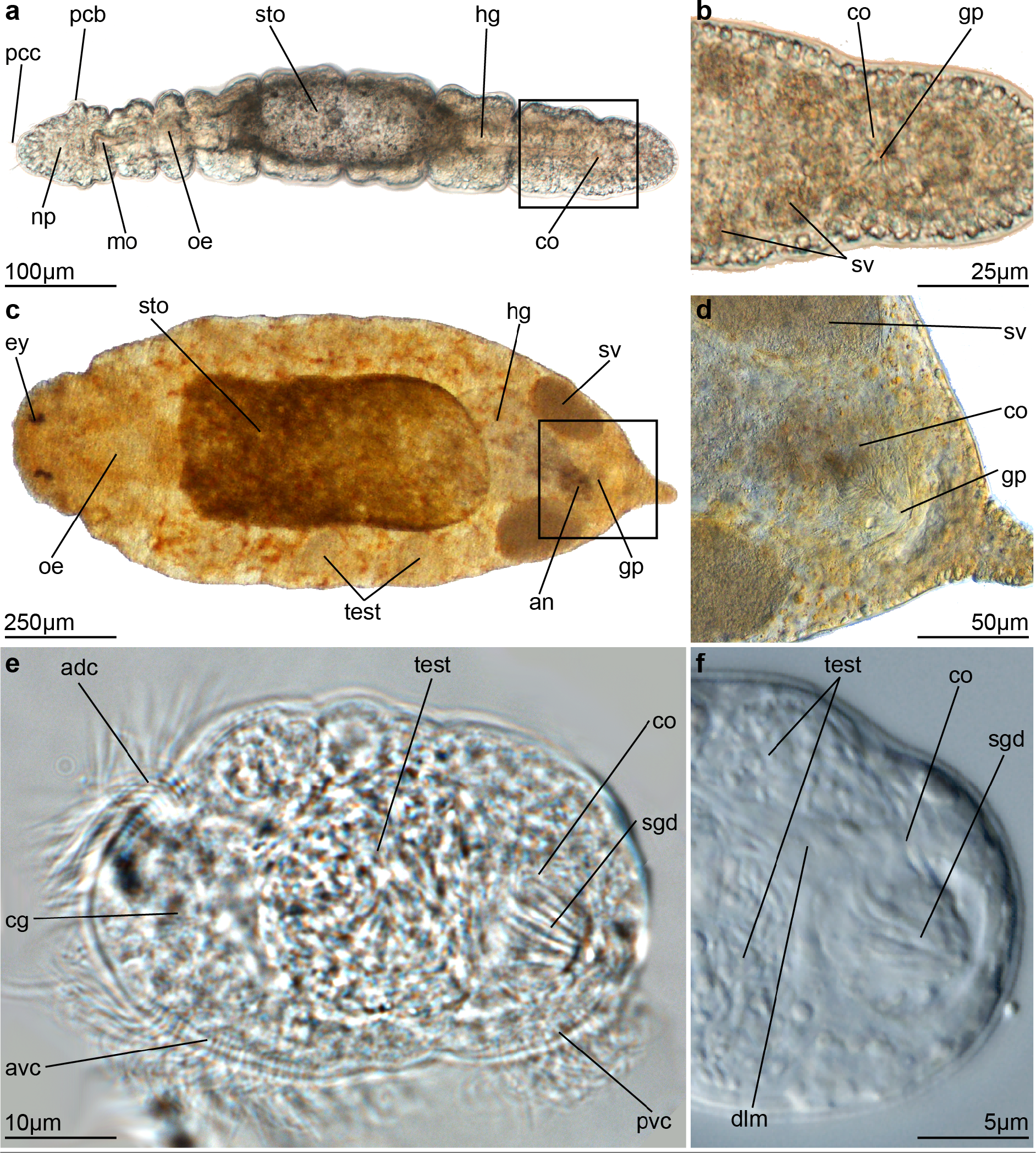
Light micrographs of adult males of *Trilobodrilus axi* (a), *Dinophilus vorticoides* (c) and *D. gyrociliatus* (e) with details of the copulatory organs (b, d, f) a, c) in dorsal view, b, d, f) ventral view, e) lateral view; anterior oriented to the left. *Abbreviations: adc – anterior dorsal ciliation, an – anus, avc – anterior ventral ciliation, cg – cerebral ganglion, co – copulatory organ (penis), dlm – dorsal longitudinal muscle, ey – eye, gp – gonopore, hg – hindgut, mo – mouth opening, np – brain neuropil, oe – oesophagus, pcb – prostomial ciliary band, pcc – prostomial compound cilia, pvc – posterior ventral ciliation, sgd – stylet gland ducts, sto – stomach, sv – seminal vesicle, test – testis*.

The paired spermioducts enter the penis bulb from each lateral side, penetrate the muscular penis sheath, and open into the lumen posterior to the penis cone. The lumen, which is also where the majority of the gland ducts seem to lead to, narrows ventrally/posteriorly towards the gonopore. The gonopore opens medially between the multiciliated epidermal cells of the ventral ciliary band. More species-specific details are described below.

The copulatory organ (penis) in all three species consists of epithelial cells alternated with gland ducts, with the glands’ cell bodies being located mainly dorsally, laterally and anteriorly to the copulatory organ (not shown). Based on previous ultrastructural and histological studies by Scharnofske [4], there are at least two different gland types involved, which differ in their vesicular or granular content. While the copulatory organ is spherical in *Trilobodrilus axi* (Figs. 1a, b, 2a, b, c, d, 4, 5), it is more elongated in *D. gyrociliatus* (1.5 times longer than wide, Figs 1e, f, 3a, b, c, d, e, 4, 5) and in *D. vorticoides* (2-3 times longer than wide, Figs 1c, d, 2e, f, g, 4, 5).

### Musculature of the male copulatory organ

The copulatory organ (penis) is located medioventrally above and between the ventral and ventrolateral longitudinal muscle bundles of the body wall (vlm, vllm, Figs 2a, b, e, f, 3a, b) with thin muscle fibres anchoring it mainly ventral to the body wall in the two larger species *D. vorticoides* and *T. axi* (am, Figs 2b, f, 4a, b), while it is directly linked to the mainly longitudinal body wall musculature in the *D. gyrociliatus* dwarf males (Figs. 3a, b, 4c). In all three species the musculature of the copulatory organ consists of an external muscular penis sheath (ps, Figs 2c, d, g, 3c, d, e, 4a, b, c) and an internal penis cone (pc, Figs 2c, d, g, 3d, e, 4a, b, c). The penis sheath is formed mainly by longitudinal muscles extending from the anterior end of the organ towards its ventral opening, where the muscles attach to the sphincter surrounding the gonopore in all three dinophilids (sph2, Figs 2c, d, f, g, 3c, d, e). In *D. gyrociliatus*, the muscle fibres are more flattened, thereby giving a more sheath-like and less organized appearance than the muscles constituting the copulatory organ of the other two studied species (see also [18,21,22] for a more detailed description). Additionally, one or two more anterior sphincter muscles surround the entire penis bulb (sph1, sph3, Figs 2c, 3b, c, d, 4a, b, c). In *T. axi*, one of them is situated approximately equatorial (where it is connected to the pair of thin longitudinal muscle bundles of the body wall musculature, sph1, Figs 2c, 4a), and an additional, thinner sphincter is found between it and the one around the gonopore (sph3, Figs. 2c, 4a). An approximately equatorial sphincter is also present in *D. vorticoides* (Figs 2e, f, g, 4b) and *D. gyrociliatus* (Figs 3c, d, 4c), although exact trajectories of muscle fibres are hard to follow in the latter due to the close proximity of these elements to each other and them being tightly intertwined. Surprisingly, the much more elongated copulatory organ of *D. vorticoides* does not necessitate additional sphincter- or circular muscles to accomplish its functions.

**Figure 2.**
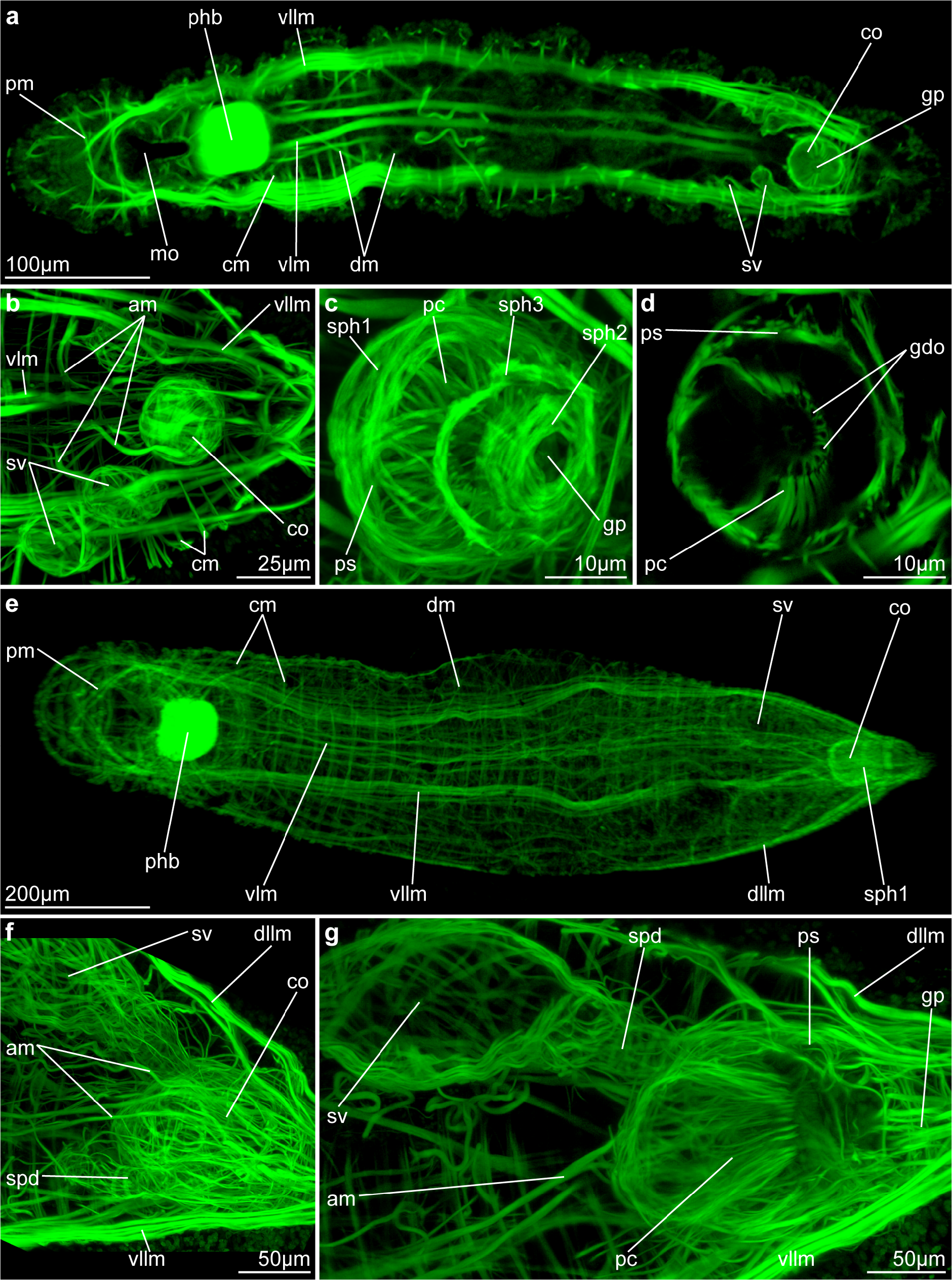
Musculature of adult males of *Trilobodrilus axi* (a-d) and *Dinophilus vorticoides* (e-g) Maximum intensity projections of CLSM-image stacks from phalloidin-treated specimens. a) Main ventral musculature in *T. axi*, ventral view, middle part of Z-stack only), b) Copulatory organ and seminal vesicles, dorsal view, ventral part of Z-stack. c) Close up of copulatory organ, ventral view. d) Horizontal section through middle part of copulatory organ showing muscles of penis sheath, penis cone and openings of gland ducts. e) Musculature of *D. vorticoides*, ventral view, ventral half of Z-stack. f) Copulatory organ and right seminal vesicle, lateral view, right half of Z-stack. g) Sagittal section through copulatory organ and right seminal vesicle, 3D crop of Z-stack. *Abbreviations: am – anchoring muscle, cm – circular muscle, co – copulatory organ (penis), dllm – dorsolateral longitudinal muscle, dm – diagonal muscle, gdo – gland duct openings, gp – gonopore, mo – mouth opening, pc – penis cone, phb – pharyngeal bulb, pm – prostomial muscle, ps – penis sheath, spd – spermioduct, sph1, 2, 3 – sphincter muscles 1, 2, 3, sv – seminal vesicle, vllm – ventrolateral longitudinal muscle, vlm – ventral longitudinal muscle*.

**Figure 3.**
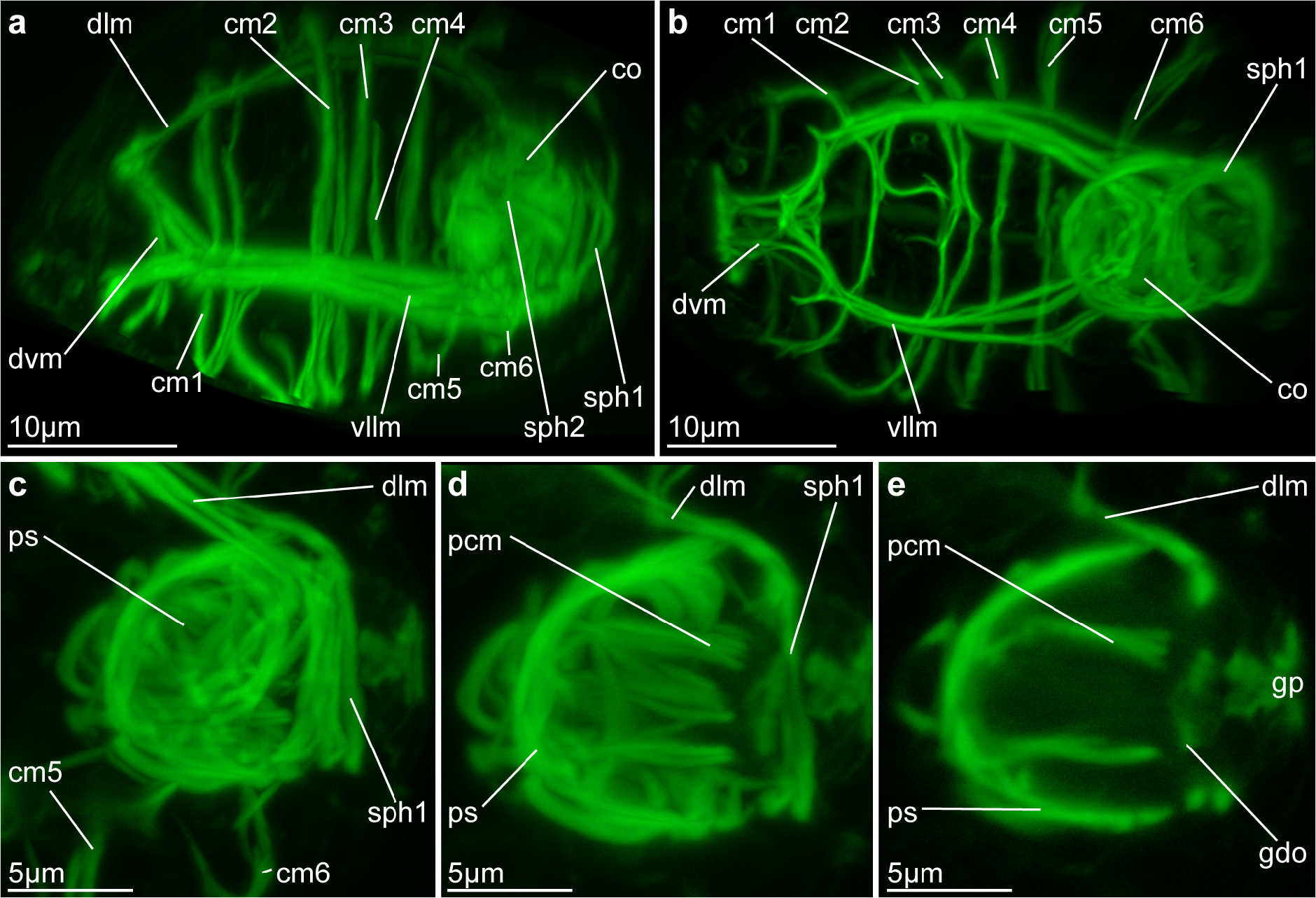
Musculature of adult dwarf males of *Dinophilus gyrociliatus*. Maximum intensity projections of CLSM-image stacks from phalloidin-treated specimens. a) Lateral and b) ventral overview, complete Z-stack. c-d) Details of copulatory organ, lateral views : c) penis sheath, 3D crop of Z-stack, d) sagittal section through lateral part of penis cone and penis sheath, e) sagittal section through middle part of penis sheath, penis cone and openings of gland ducts. *Abbreviations: cm1-6 – circular muscle 1-6, co – copulatory organ (penis), dlm – dorsal longitudinal muscle, dvm – dorsoventral muscle, gdo – gland duct openings, gp gonopore, pcm – penis cone muscle, ps – penis sheath, sph1,2 – sphincter muscle 1,2, vllm ventrolateral longitudinal muscle*.

The inner penis cone is in all three species constituted by a thin layer of longitudinal muscle fibres surrounding and supporting the internal gland ducts, whose cell bodies are mostly located outside the penis bulb [4,5,6,21,22]. The exclusively longitudinal muscle fibres of the penis cone originate at the anterior end of the copulatory organ (antero-dorsally in *T. axi*), follow the penis sheath internally for approximately half the length of the penis cone, and then deviate inwards (Figs 2d, g, 3d, e, 4a, b, c). In *T. axi*, the penis cone is located more antero-dorsally and cannot reach the posterior gonopore or even get protruded through it. According to [4], the glands whose ducts line the posterior part of the penis internally belong to a different cell type than the ones in its anterior part, or the ones of the penis sheath. The glands, whose openings surround the gonopore, most likely aid to attachment during copulation [4] similar to the adhesive glands reported near the gonopore opening in *D. vorticoides* [5] and *D. gyrociliatus* [21]. In *D. vorticoides* and *D. gyrociliatus*, the penis cone extends more posteriorly (Figs 2g, 3d, e, 4b, c) with stylet glands forming the tapering end of the penis cone directed towards the gonopore. According to histological investigations [5], the males use the adhesive glands, which are located inside the penis sheath in close proximity to the gonopore, to attach themselves to the females, while the glands, whose ducts are leading into the penis cone, produce the secrete to enzymatically dissolve the epidermis. The stylet glands, or more precisely their rod-shaped secretions, in *Dinophilus*-species attach to the edges of the thus created hole in the epidermis and thereby keep the previously enzymatically dissolved hole open. Jägersten [5] described them to only hold on to the edges while the penis cone retracts in *D. vorticoides* (in his description *D.* cf. *taeniatus*), and then aid their fastening to a second, more internally located population of adhesive glands. However, neither previous nor this study found similar adhesive glands in *D. gyrociliatus* or *T. axi* (see also [4,8,18,21,22]. In all studied species sperm (transported into the lumen of the penis sheath by the laterally opening spermioducts) is transported into the female through the gonopore by a mixture of ciliary movement of the spermioducts and muscular pumping of the penis [4,5,7,8] after an initial retraction of the penis cone. The movement is possibly aided by contractions of the body wall musculature and the pressure of the filled testes in especially the miniaturized *D. gyrociliatus*.

The seminal vesicles are situated anterior and dorsolateral to the copulatory organ in *T. axi* (sv, Figs 2a, b, 4a) and *D. vorticoides* (Figs 2e, f, g, 4b), and are surrounded by a thin layer of longitudinal and diagonal muscle fibres with thin muscles anchoring them to the lateral body wall. Thin and innervated (see below) sphincter muscles at the opening passage between the testes and seminal vesicles are only found in *T. axi* (Figs 4a, d, 5a, d). The spermioducts of the other two species are lined by few longitudinal and several, quite widely spaced circular muscles (spd, Figs 2f, g, 4a, b).

**Figure 4.**
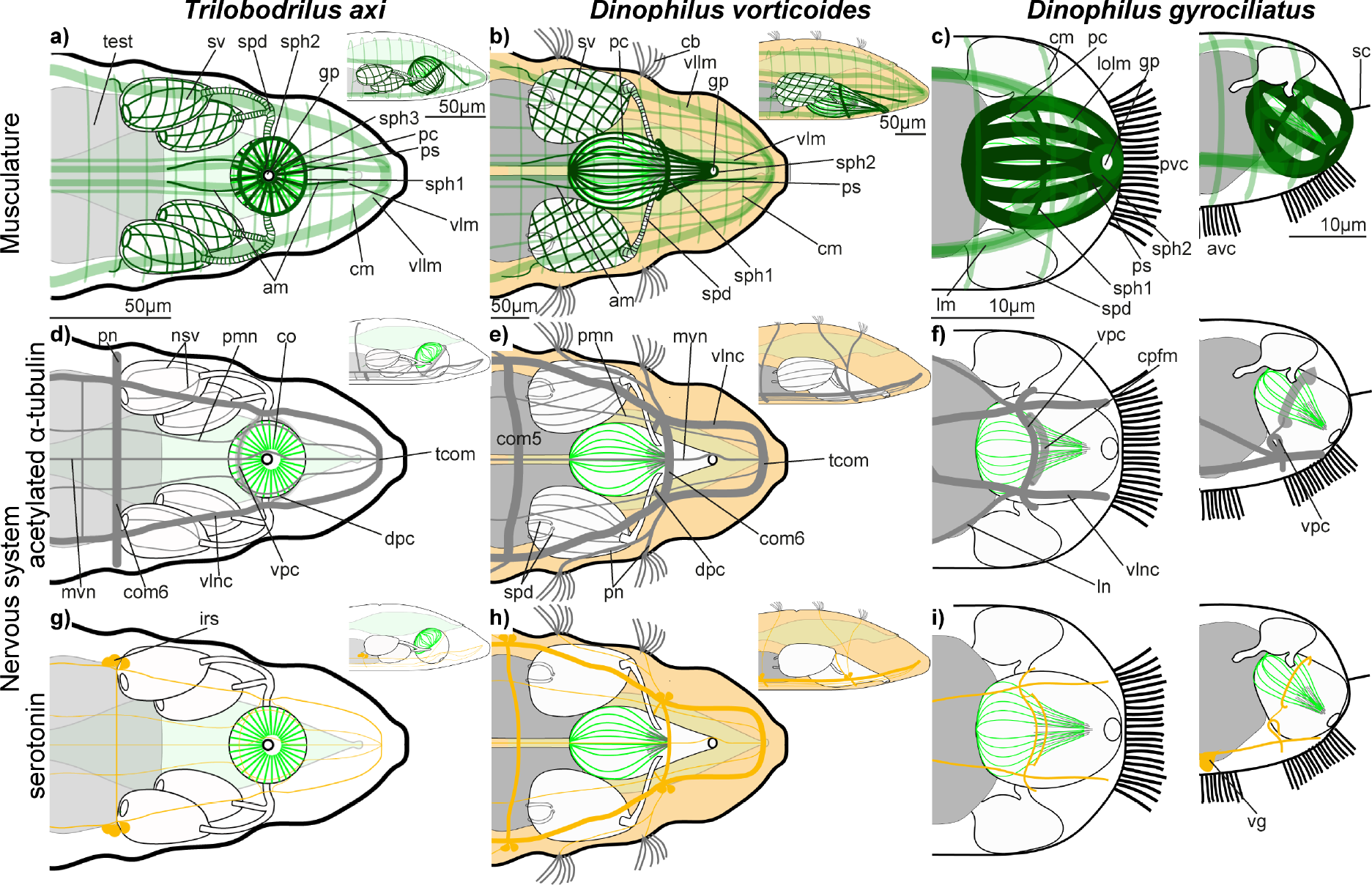
Schematic reconstruction of phalloidin stained musculature (a-c) and nervous system of the copulatory organ as revealed by acetylated α-tubulin immunoreactivity (d-f) and serotonin-like immunoreactivity (g-i) of male Dinophilidae, based on CLSM Z-stacks. These drawings represent the morphology of male *Trilobodrilus axi* (a, d, g), *Dinophilus vorticoides* (b, e, h) and *D. gyrociliatus* (c, f, i) in ventral (large drawing), and lateral view (small drawing). Larger drawings only showing ventral half of animal (all dorsal muscles and nerves omitted, except for *D. gyrociliatus*, whose reproductive system extends further dorsally). The musculature of the *D. gyrociliatus* copulatory organ is highly simplified, for more details see [18,21,22]. Thin black lines in the tip of the penis cone in *D. vorticoides* and *D. gyrociliatus* represent the rod-like secretes of the stylet glands. Body wall muscles in a-c) are shown more hyaline to allow for a better presentation of the muscles of the reproductive system. *Abbreviations: am – anchor muscles, avc – anterior ventral ciliation, cb – ciliary band, cm – circular muscle, co – copulatory organ (penis), com5, 6 – fifth/sixth body commissure, cpfm – circumpenial fibre mass, dpc – dorsal penial commissure, gp – gonopore, lm – longitudinal muscle of the body wall, ln – laternal nerve, lolm – loops of the penis sheath formed by the longitudinal body wall muscles, mvn – medioventral nerve, nsv – nerves of the seminal vesicles, pc – penis cone, pcom – penial commissure, pmn – paramedian nerve, pn – peripheral nerve, ps – penis sheath, pvc – posterior ventral ciliation, sc – sensory cilium, spd – spermioduct, sph – sphincter, sv – seminal vesicle, tcom – terminal commissure, test – testis, vg – ventral ganglion, vllm – ventrolateral longitudinal nerve, vlm – ventral longitudinal muscle(s), vlnc – ventrolateral nerve cord, vpc – ventral penial commissure*.

No musculature was found around the testes in any species.

### Neural innervation of the male copulatory organ

The architecture of the ventral nervous system of the here used three dinophilid species is described in detail in previous studies [10,11,18,21,22]. In the two “normal-sized” males of the monomorphic *T. axi* and *D. vorticoides* the central nervous system comprises the brain, circumesophageal connectives, two main ventrolateral nerve cords, two paramedian longitudinal nerves and one unpaired midventral longitudinal nerve. The two main ventrolateral nerve cords originate in the brain and extend throughout the body external to the ventrolateral longitudinal muscle bundles and fuse with the terminal commissure, which is located ventroposterior to the anus. The pair of paramedian and the unpaired medioventral longitudinal nerves extend posteriorly from their origin at the subesophageal commissure posterior to the mouth opening. The paramedian nerves deflect laterally around the copulatory organ, and subsequently follow the ventrolateral nerve cords closely, before fusing with the ventrolateral nerve cords and the medioventral nerve at the terminal commissure. All longitudinal nerves are connected by commissures in each segment. While *D. vorticoides* has one commissure in the sixth (most posterior) segment, two to three commissures of different thickness (i.e. different number of neurites) [10,11,32] are found in *T. axi*, with the most prominent commissure here termed the main commissure (Figs Figs 4, 5, 6, 7a, e,). *Dinophilus gyrociliatus* dwarf males show a much more reduced nervous system, with one pair of ventral and one pair of lateral nerve cords, each consisting of only few neurites and connected to each other by the cerebral, ventral and penial commissures (Figs 4, 5, 7a, [21,22,32])

**Figure 5.**
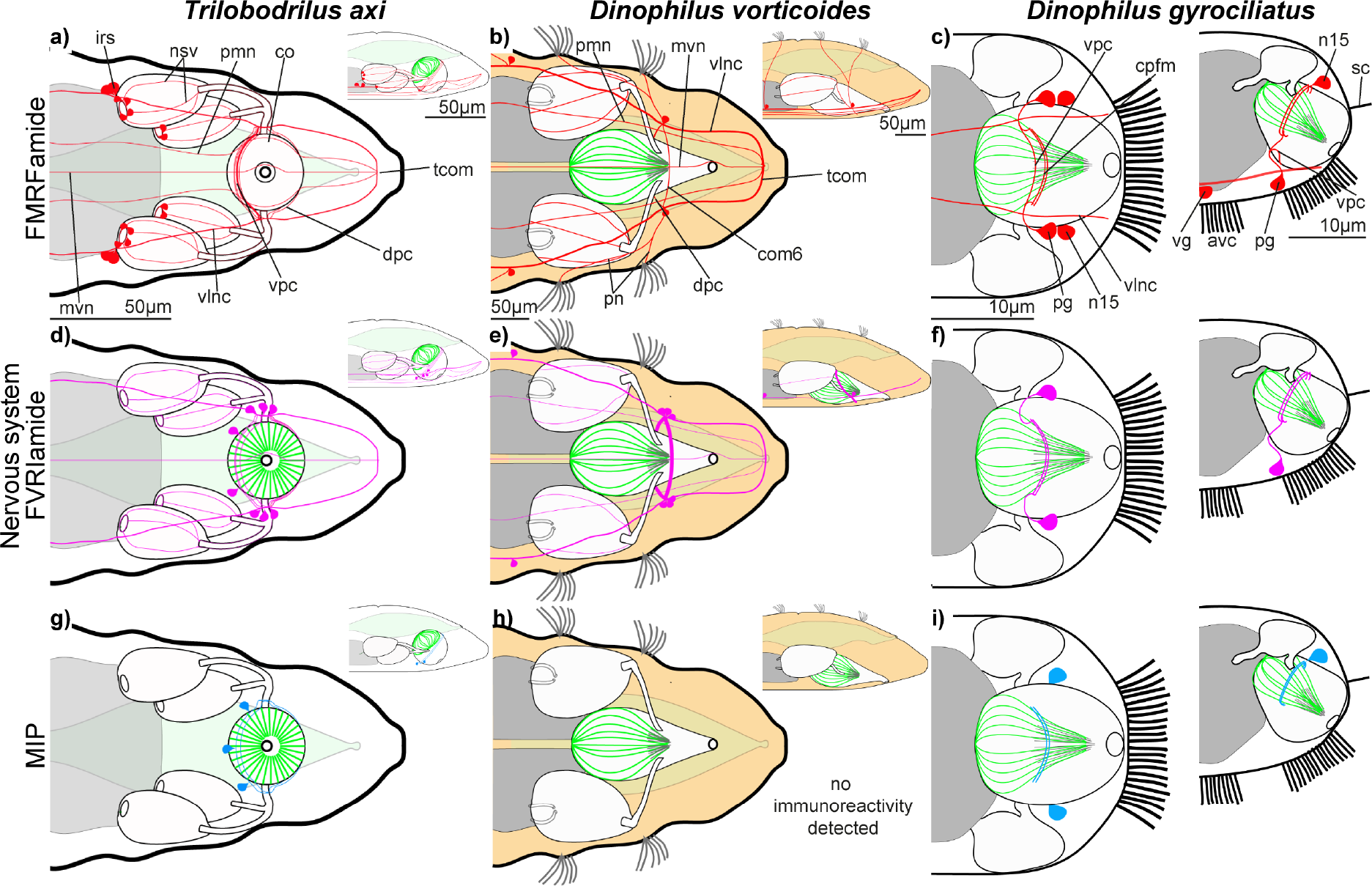
Schematic reconstruction of the nervous system of the copulatory organ as revealed by FMRFamide-like (a-c), FVRIamide-like (d-f), and MIP-like immunoreactivity (g, h) of male Dinophilidae, based on CLSM Z-stacks. These drawings represent the morphology of the males of *Trilobodrilus axi* (a, d, g), *Dinophilus vorticoides* (b, e) and *D. gyrociliatus* (c, f, h) in ventral (large drawing), and lateral view (small inset). In the larger drawing only the ventral half of the animal is shown (all dorsal nerves are omitted, except in *D. gyrociliatus*, whose reproductive system extends further dorsally). Thin black lines in the tip of the penis cone in *D. vorticoides* and *D. gyrociliatus* represent the rod-like secretes of the stylet glands. *Abbreviations: co – copulatory organ (penis), com6 – sixth body commissure, cpfm – circumpenial fibre mass, dpc – dorsal penial commissure, mvn – medioventral nerve, nsv – nerves of the seminal vesicles, n15 – neuron 15 (nomenclature based on Windoffer & Westheide 1988a, b), pcom – penial commissure, pg – penial ganglion, pmn – paramedian nerve, pn – peripheral nerve, tcom – terminal commissure, vg – ventral ganglion, vlnc – ventrolateral nerve cord, vpc – ventral penial commissure*.

**Figure 6.**
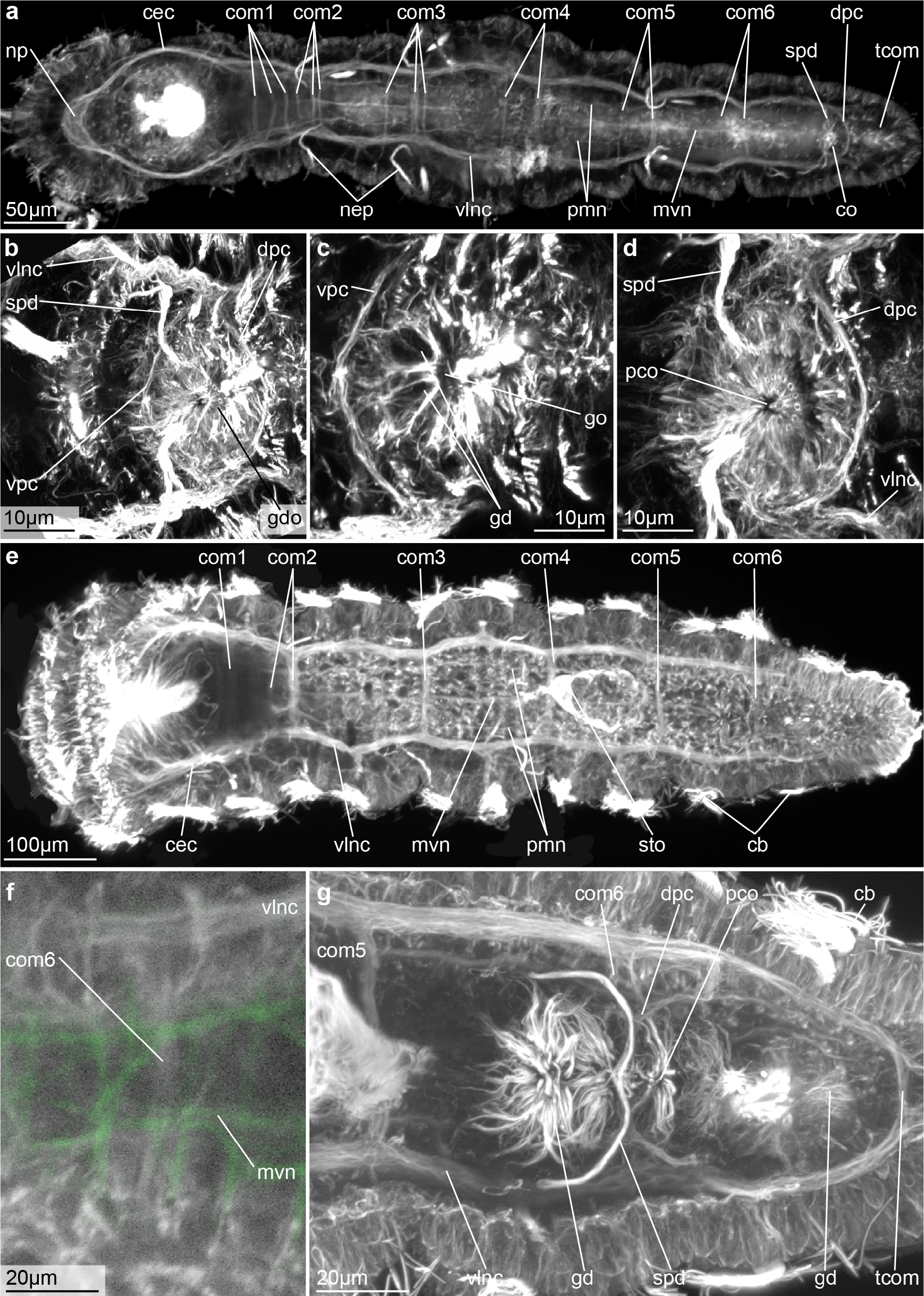
Nervous system of adult males and their copulatory organs in *Trilobodrilus axi* (a-d) and *Dinophilus vorticoides* (e-g), as shown from acetylated α-tubulin immunoreactivity. Maximum intensity projections of CLSM-images. a) Ventral overview of the nervous system of *T. axi*, 3D crop of ventral part of Z-stack. b-d) Nervous system around copulatory organ, showing the ventral penial commissure (b, c), and the dorsal penial commissure (d). e) Ventral overview of the nervous system of *D. vorticoides*, 3D crop of ventral part of Z-stack. f) Close up showing the commissure of the sixth body segment (potentially fused with the vpc?), ventral view, 3D crop of ventral part of Z-stack. g) Innervation of the copulatory organ, ventral view, ventro-median part of Z-stack. *Abbreviations: cb – ciliary band, cec – circumesophageal connective, co – copulatory organ (penis), com1-6 – commissures of the body segments 1-6, dpc – dorsal penial commissure, gd glandular ducts, gdo – glandular duct openings, go – gonopore, mvn – medioventral nerve, nep – nephridium, np – brain neuropil, pco – penis cone opening, pmn – paramedian nerve, spd – spermioduct, sto – stomach, tcom – terminal commissure, vlnc – ventrolateral nerve cord*.

**Figure 7.**
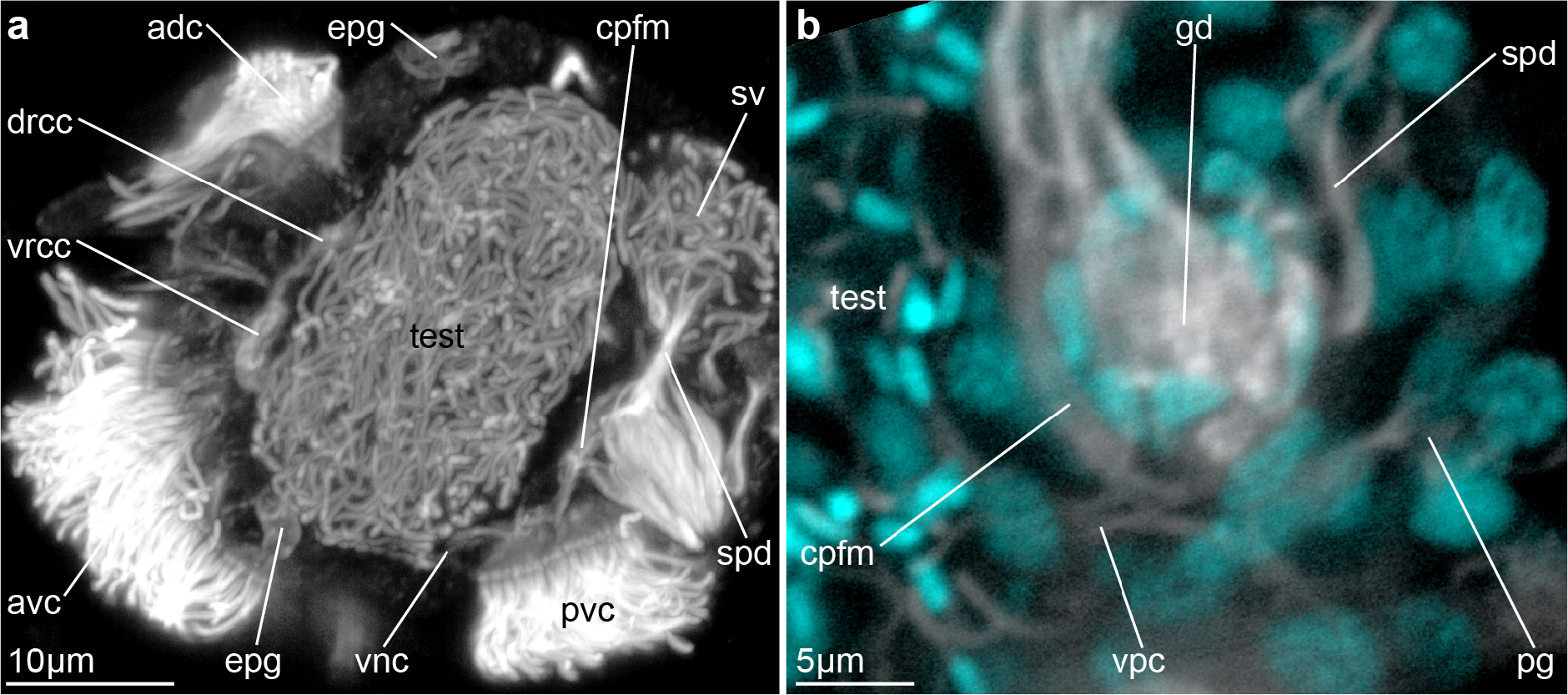
Nervous system and copulatory organ of adult *Dinophilus gyrociliatus* dwarf males as shown from acetylated α-tubulin immunoreactivity. Maximum intensity projections of CLSM-images. a) Lateral overview of a dwarf male (complete Z-stack), and b) posterior transverse section through the copulatory organ (also showing nuclei with DAPI-4 staining in cyan). *Abbreviations: adc – anterior dorsal ciliation, avc – anterior ventral ciliation, cpfm – circumpenial fibre mass, drcc – dorsal root of the circumesophageal connective, epg – epidermal gland, gd – gland ducts, pg – penis ganglion, pvc – posterior ventral commissure, spd – spermioduct, sv – seminal vesicle, test – testis, vnc – ventral nerve cord, vrcc – ventral root of the circumesophageal commissure*.

In contrast to the high degree of similarities in the musculature and the overall nervous system of the three dinophilid species, the detailed neural innervation of the copulatory organ differs slightly among them: male *T. axi* possess one horizontal, slightly oblique nerve ring around the penis tracing the median sphincter muscle. This circumpenial fibre mass is partly composed of two commissures originating at slightly different points of the ventral cords and embracing the copulatory organ ventrally and dorsally, respectively, with the ventral commissure being more condensed and anterior to the dorsal one (Figs 4d, 6b, c, d, 7a, d, g, j). Male *D. vorticoides* also have a ventral and a dorsal commissure; however, here the dorsal commissure is located anterior to the ventral commissure (Figs 4e, 6f, g, 8b, e, h). Furthermore, the ventral penial commissure is either formed by or fused with the ventral commissure of the sixth body segment (com6), whereas the ventral penial commissure is a separate, independent formation in *T. axi* posterior to the commissures of the sixth body segment (compare Figs 4d, 6a, e). Yet another architecture is found in the dwarf male of *D. gyrociliatus* (Figs 4, 5, 7b, [21,22]): here, a commissure is formed between the ganglia ventral to the copulatory organ (maybe corresponding to the ventral penial commissure in *D. vorticoides* and *T. axi,* Figs 4d, f, 5, 7b), and a ring-shaped circumpenial fibre mass is formed around the penis cone within the penis sheath (Figs 4g, i, 5c, f, h, 7a, b, 8c, f, I, k), which seems to be different to the other, externally to the muscular penis sheath formed neural structures in *D. vorticoides* and *T. axi* [4,5,21]. Yet, the penis sheath in *D. gyrociliatus* is formed by much fewer muscle cells and additionally joined by body wall muscles [18], which according to Windoffer & Westheide [21] may be folded within each other and possibly also around the circumpenial fibre mass, explaining its deviating position within the penis sheath.

Neural innervation of the seminal vesicles could be traced in the two monomorphic species *D. vorticoides* and *T. axi*, where individual nerve fibres extend from the circumpenial nerve ring along the spermioducts and the seminal vesicles. In *T. axi*, they terminate at the nerve ring surrounding the testes openings (and sphincter muscles), in *D. vorticoides* we could not find the termination of the longitudinal fibres. These fine nerves are weakly labelled by acetylated α-tubulin immunoreactivity, but better visible with the specific neurotransmitter immunoreactivities of FMRFamide and FVRIamide (see below). The neural innervation of the spermioducts and seminal vesicles in *D. gyrociliatus* is much weaker and consists of only one to two neurites, which extend towards the testes (Fig. 4, raw data not shown).

### Neurotransmitter immunoreactivity patterns

Serotonin-like immunoreactive (serotonin-LIR) fibres are found in nearly all parts of the nervous system e.g. in the brain neuropil, ventrolateral nerve cords, paramedian and median nerves as well as some of the commissures and their associated serotonin-LIR somata (Figs 4g, h, i, 6a, [10,11,14]). Likewise, the ventral and dorsal penial commissures in *T. axi* (Figs 4, 8a, Table 1) and *D. vorticoides* contain serotonin-LIR fibres (Figs 4, 8b, Table 1) as well as the ventral commissure and the circumpenial fibre mass (with stronger immunoreactivity in the dorsal region of the fibre mass) in *D. gyrociliatus* (Figs 4, 8c, Table 1). There furthermore is one pair of immunoreactive somata in close proximity to the nervous system around the copulatory organs of *D. vorticoides* and *D. gyrociliatus*, though we could not trace these somata’s relation to the immunoreactive neurites of the penial innervation. It seems that the labelled fibres mainly originate from the somata along the ventrolateral nerve cords, and not from somata, which are close to the penial commissures or the circumpenial fibre mass. Serotonin-LIR was not detected in the nerves running along the seminal vesicles and spermioducts (Figs 4g, h, i, 8a, b).

**Table 1.**
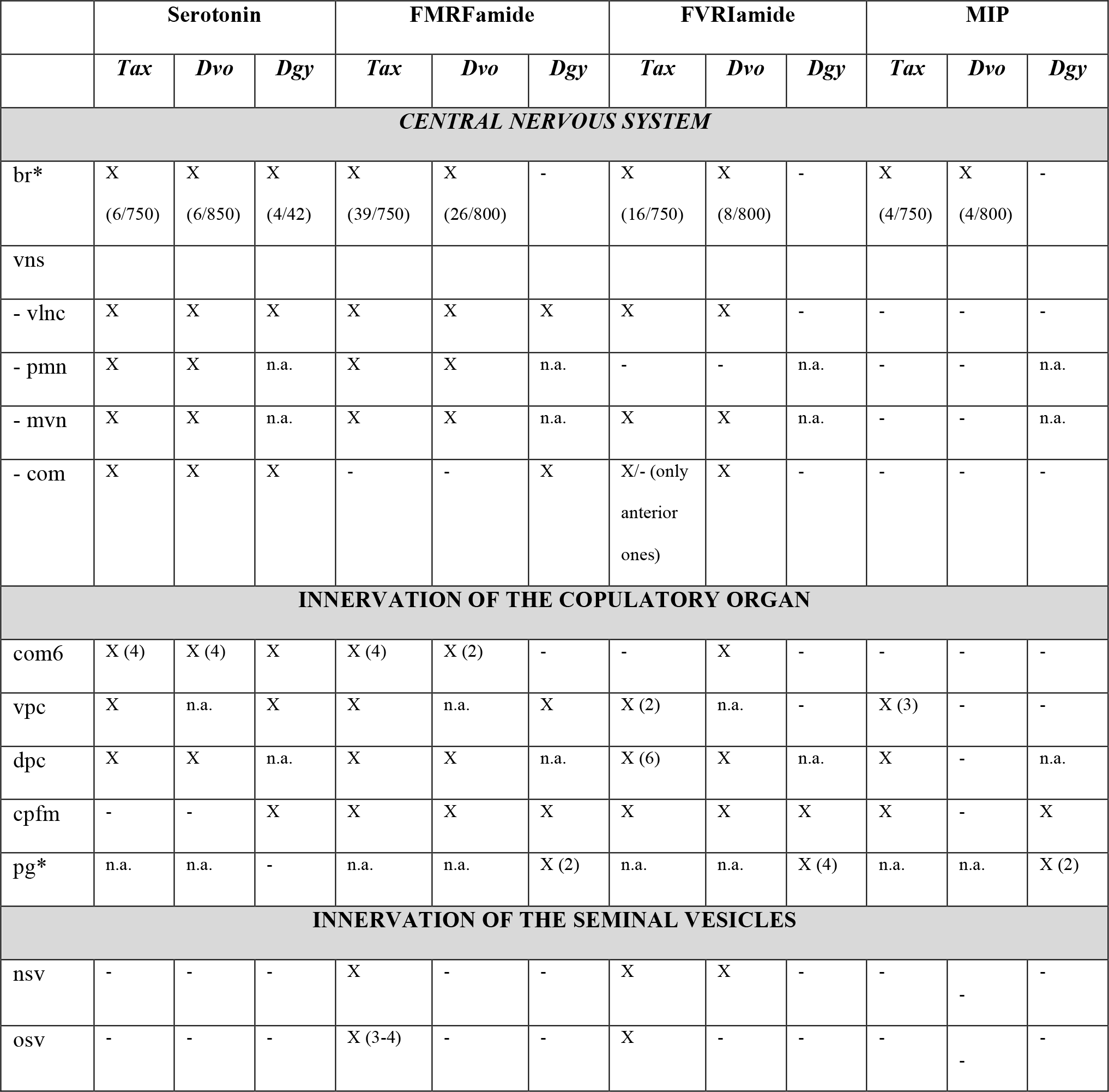
Immunoreactivity patterns in the nervous system possibly innervating the copulatory organ of *Trilobodrilus axi* (Tax), *Dinophilus vorticoides* (Dvo) and *D. gyrociliatus* (Dgy) Positive immunoreactive is indicated by “X”, with the number of immunoreactive perikarya indicated by the number within parentheses. For the numbers of immunoreactive perikarya in the brain and the penial ganglia, their number is given next to the total number of perikarya of the ganglia. “−“ indicates the absence of immunoreactive signal, “n.a.” (not applicable) is used if a neural structure is not present in the respective species. Due to our findings and previous reports, we argue for the possible homology of the sixth body commissure (“com6”) in *T. axi* and *D. vorticoides* and the ventral penial commissure between the two penial ganglia in *D. gyrociliatus*. We furthermore argue that the circumpenial fibre mass in *D. gyrociliatus* is possibly homologous to the nerve ring around the copulatory organ in *T. axi* and possibly also the ventral and dorsal penial commissures in *vorticoides* and *T. axi*, however, we kept the latter separate in the table below to facilitate our comparison. In *Dinophilus gyrociliatus* dwarf males the sensory cells associated to the cerebral or penial ganglia are included in the number of brain or penial ganglion cells, which is indicated by “*”. *Abbreviations: br – brain, com – commissure, com6 – commissure 6 (in* D. gyrociliatus*, this refers to the commissure between the ventral ganglia), cpfm – circumpenial fibre mass, dpc – dorsal penial commissure, mvn – medioventral nerve, nsv - longitudinal nerve fibres along the seminal vesicles, osv – nerve ring around the opening of the seminal vesicles, pcom – penis commissure (in* D. gyrociliatus*), pg – penial ganglion, pmn – paramedian nerve, vlnc – ventrolateral nerve cord, vns – ventral nervous system, vpc – ventral penial commissure*.

**Figure 8.**
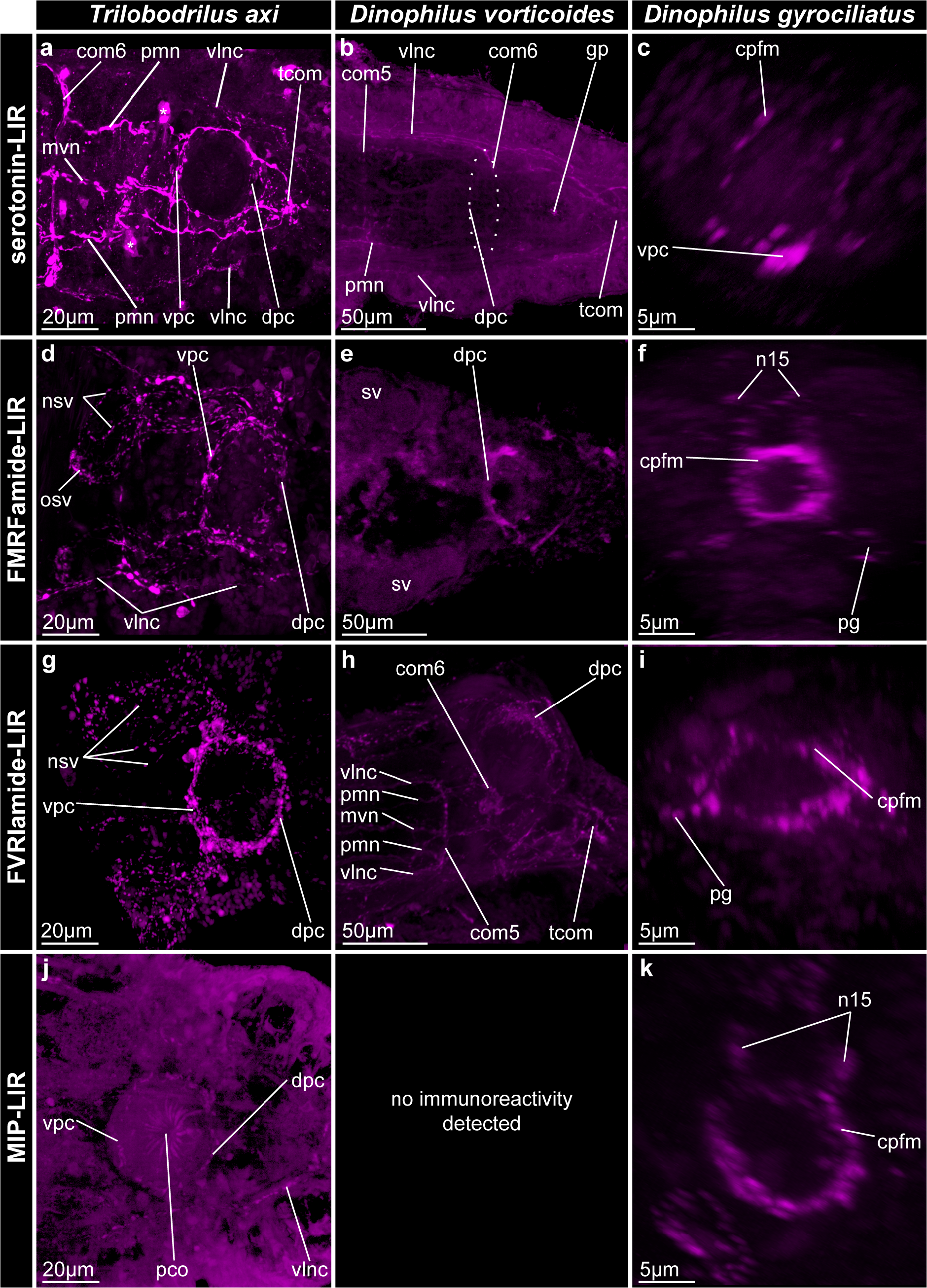
Immunoreactivity patterns of serotonin (a-c), FMRFamide (d-f), FVRIamide (g-i), and MIP (j, k) in the nervous system of the copulatory organ in adult males of *Trilobodrilus axi* (a, d, g, j), *Dinophilus vorticoides* (b, e, h), and *D. gyrociliatus* (c, f, I, k) Maximum intensity projections of CLSM-images. a, b, d, e, g, h, j) Ventral view, ventral part of Z-stack, c, f, i, k) posterior view, transverse 3D crop through the copulatory organ. For the large *D. vorticoides* (b, e, h) only a small subsample of the entire stack has been compiled into a 2D image, omitting some of the traits described in Figs 4 and 5, but see [11, 18] for more information on the overall nervous system. The asterisks in (a) mark artefacts (a fold in the animal led to a stronger accumulation of serotonin-LIR) and no real serotonin-LIR of neural elements. *Abbreviations: com5, 6 – commissure of the fifth/sixth body segment, cpfm – circumpenial fibre mass, dpc – dorsal penial commissure, gd – gland ducts, gp – gonopore, mvn – medioventral nerve, nsv – nerves of the seminal vesicles, n15 – neurite extending from dorsoposterior sensory neuron 15 (nomenclature based on Windoffer & Westheide 1988a, b), osv – opening of the seminal vesicles to the testes, pc – penis cone, pco – penis cone opening, pg – penial gamglion, pmn – paramedian nerve, pvc – posterior ventral ciliation, sv – seminal vesicle, test – test, vlnc – ventrolateral nerve cord, vpc – ventral penial commissure*.

FMRFamide-LIR is equally abundant to serotonin-LIR in the overall nervous system of males of *T. axi* and *D. vorticoides*, but more restricted to the posterior region of the body in in the dwarf males of *D. gyrociliatus*. FMRFamide-LIR fibres are thus found in all commissures around and within the copulatory organ (ventral and dorsal penial commissures of *T. axi* (Figs 5a, 8d, Table 1) and *D. vorticoides* (Figs 5b, 8e, Table 1), and penial commissure and circumpenial fibre mass in *D. gyrociliatus* (Figs 5c, 8e, Table 1)). Several nerves around the seminal vesicles in *T. axi* (lnsv, Figs 5a, 8e) and *D. vorticoides* (Figs 5b, raw data not shown) exhibit FMRFamide-like immunoreactivity, and male *T. axi* possess additional FMRFamide-LIR somata around the openings to the testes (osv, Figs 5a, 8d). No FMRFamide-LIR nerves are lining the spermioducts of *D. gyrociliatus*, but one pair of cells of the penial ganglion and one pair of cells dorsal to the circumpenial fibre mass exhibit this immunoreactivity (Fig 5c).

FVRIamide-LIR fibres and somata are present in the brain neuropil and in the ventral nervous system of the first three segments, but are most abundant in the stomatogastric nervous system in the middle to posterior region of the body (data not shown here, but see [11] for more details). FVRIamide-like immunoreactivity is furthermore detected around the copulatory organ in *T. axi* and *D. vorticoides*: eight (in *T. axi*) or four (in *D. vorticoides*) immunoreactive somata are arranged around the penis, and project immunoreactive fibres longitudinally along the spermioducts and the seminal vesicles (Figs 5d, e, 8g, h, Table 1).

Additionally, FVRI-LIR fibres without associated immunoreactive somata are forming a ring around the opening of the testes to the seminal vesicles in *T. axi*, probably tracing the sphincter muscle and regulating the transfer of sperm to the seminal vesicles (data not shown). In *D. gyrociliatus* dwarf males FVRIamide seems to be restricted to the copulatory organ and the associated neurons in the posterior region of the body (Figs 5f, 8i, Table 1). Here, it is located in the circumpenial fibre mass (Figs 5f, 8i) and one pair of cells in the penial ganglia.

Myoinhibitory peptide (MIP)-like immunoreactivity only shows a weak signal in all studied specimens, thereby complicating its exact localization in the nervous system. In *T. axi* one to two immunoreactive fibres are found in the anterior part of the ventrolateral nerve cords (but not in the paramedian or medioventral nerves) supplemented by one pair of immunoreactive somata and one to two MIP-LIR neurites in the main commissure of the third segment. No other segmental commissures were found to comprise MIP-like immunoreactivity [11]. Yet, several specimens of *T. axi* show three immunoreactive somata anteriomedian to the copulatory organ; one of them anterior, and the other two located lateral on each side of the spherical penis (Fig 5g, Table 1). These somata are connected to each other by a few MIP-LIR fibres, which are part of the circumpenial fibre mass around the copulatory organ (Figs 5g, 8j). No MIP-like immunoreactivity is detected in neither fibres nor somata associated with the seminal vesicles and spermioducts. No data about the distribution of MIP-like immunoreactivity in the nervous system around the reproductive organs was obtained despite several attempts for *D. vorticoides*. In dwarf males of *D. gyrociliatus*, MIP-LIR fibres are found solely in the circumpenial fibre mass (two to three fibres, Figs 5i, 8k, Table 1), and potentially in the somata of dorsal sensory neurons, however, the latter pattern is extremely weak and it was not possible to relocate these cells in all tested animals (Fig 5i).

## Discussion

### Muscular similarities in dinophilid copulatory organs

Using phalloidin-labelling and CLSM, this study succeeded in revealing the three-dimensional structure of the reproductive organs in three dinophilid species, with special focus on the copulatory organ and its proposed function. Despite the differences in size and life strategy of the three species as well as of the different overall shape of the copulatory organ (elongated to pear-shaped and oriented along the anterior-posterior axis in *D. vorticoides* (Figs 1c, d, 2e, f, g) and *D. gyrociliatus* (Figs 1e, f, 2a, b, c, d, e), spherical and oriented along the dorsoventral body axis in *T. axi* (Figs 1a, b, 2a, b, c, d)), the copulatory organ in all species comprises an external muscular penis sheath and an internal penis cone (Figs 2, 3). The penis sheath consists of circular muscles (forming one or two sphincters) external to its less organized, but mainly longitudinally oriented muscle layer [4,5,7]. The penis cone is located within the anterior part of the penis sheath (and anchored to it) and comprises only thin, longitudinal muscle fibres. In *D. vorticoides* and *D. gyrociliatus* these thin fibres trace the stylet gland ducts and in *T. axi* they form a thin and almost transverse layer in the middle part of its spherical copulatory organ, where they bend inwards. The muscles of the penis cone (also due to the glands associated with it) supposedly regulate the release of glandular products from the penis cone into the penis sheath, and from there to the gonopore for dissolving the female’s epidermis [4,5], and are also involved in the transport of sperm from the dwarf male into the female to some extent. The highly muscular penis sheath is involved in the main pumping movements of the penis, as well as in the intricate regulation of the attachment of the dwarf male to the female by orienting the adhesive glands accordingly. In contrast to previous findings about the copulatory organ being a possibly new formation in the otherwise reduced body organization [18], our comparative approach reveals the complex, two-layered penis to be a conserved trait of the annelid family Dinophilidae, which does not seem to be strongly affected by changes in size or life cycle of the species.

### Function of the dinophilid penis muscles

Most of what we know about the function of the individual muscle layers of the copulatory organ and their interactions with the different glandular and supporting cells is based on light microscopic observations and previously published, mainly morphological studies [4,5,8]. Combining this study’s results with previous histochemical analyses [4,5] and life observations [8] of *Dinophilus*, we suggest that the compartmentalization of the copulatory organ into penis sheath and penis cone serves also as a spatial separation of glands with different content and probably also different enzymatic activities. Studies focussing on the Swedish form of *D. vorticoides* have suggested that the glands in the penis cone mainly serve the enzymatic disruption of the female’s epidermis, which is accomplished by the cone’s protrusion after the adhesive glands of the penis sheath have attached the male gonopore to the female epidermis [5]. This seems to also be the case in the other two dinophilid species *D. gyrociliatus* and *T. axi* with behavioural and ultrastructural studies [4,8]. The major part of the protrusion of the penis cone is facilitated by contractions of the penis sheath. Additionally, a contraction of the thin fibres of the penis cone probably causes a widening of the penis cone opening (and thereby an increased flow of enzymes into the lumen of the penis sheath and thereby onto the female’s epidermis). Once the enzymes created the hole in the female epidermis, the penis cone is retracted, which is most likely accomplished by both retractions of the muscle fibres of the penis cone and contractions of the circular muscles of the penis sheath, leading to an overall longitudinal extension of the sheath. The retraction of the penis cone also frees passage for the sperm, which flows from the testes via the spermioducts into the lumen of the penis sheath. Pumping motions of the penis sheath and cone alternately pump sperm into the female [5,8], and are further aided by body contractions (especially in the microscopic dwarf male of *D. gyrociliatus*) and the musculature of seminal vesicles and spermioducts, ciliation, and possibly also the pressure within the fully filled testes. Due to the proposed and occasionally observed fast healing properties in these animals the hole in the female closes soon again after being released from the male copulatory organ, possibly supported by enzymes produced by the males and secreted into the open wound [4,5,8]. Our preliminary behavioural observations as well as previously published studies suggest a largely similar process in both species of *Dinophilus*, however, we have less information on *T. axi*, which differs in lacking stylet glands. This lack is possibly compensated by *Trilobodrilus* having a somewhat extended courtship behaviour, where the male and the female are crawling alongside each other for a while with occasional sensing tail sweeps of the male prior to attachment and copulation [5,7,8,20,33,34], maybe rendering the adhesion provided by the stylet glands less needed.

### Homologous orchestration of copulation in dinophilids?

The similar muscle layout of the copulatory organ is not reflected by an entirely similar neural innervation in the three studied species, although in all three species we depict a potentially similar overall function in controlling the pumping motion of the copulatory organ and transfer of sperm. This inferred functional similarity is strongest supported by an overall similar construct of a prominent nerve ring around the copulatory organ (either continuous or constituted by a ventral and a dorsal commissure between the pair of ventrolateral nerve cords), which is observed in all three species with immunoreactivities against the monoamine neurotransmitter serotonin (Figs 4g, h, I, 8a, c), as well as the neuropeptides FMRFamide (Figs 5a, b, c, 8d, e, f), FVRIamide (Figs 5d, e, f, 8g, h, i) and in *T. axi* and *D. gyrociliatus* with MIP (Figs 5g, h, 8j, k, Table 1). Acetylated α-tubulin immunoreactivity better reveals the condensed ventral and dorsal commissures in *T. axi* and *D. vorticoides* (Figs 4d, e, 6b, c, d, f, g).

Yet, the position of the circumpenial fibre mass in *D. gyrociliatus* dwarf males between the penis cone and penis sheath differs from the location of the circumpenial nerve ring external to the penis sheath in *T. axi* and *D. vorticoides*. This however might be explained by development and detailed morphology of the copulatory organ: the longitudinal body wall musculature in *D. gyrociliatus* is strongly involved in the formation of the muscle layer of the penis sheath [18], which is not the case in *T. axi* and *D. vorticoides*.

The location of the ventral commissure of the copulatory organ vis-à-vis the segmental commissures of the body seems to differ slightly between the three species: In male *T. axi* it is formed anterior to the copulatory organ and between the sixth body commissure and the terminal commissure (Figs 4d, 6b, c). In males of *D. vorticoides* it seems to either be fused with the sixth body commissure or to be replaced by it, but developmental studies are needed to fully clarify this (Figs 4e, 6e, f). The dwarf males of *D. gyrociliatus* show a ventral commissure (ventral to the circumpenial fibre mass) between the two penial ganglia (Figs 4f, 7b), which constitutes the third and posteriormost commissure besides the brain and ventral ganglia commissures. In general, the ventral penial commissure (maybe as part of the circumpenial nerve ring) seems to be an additional formation to the ventral cord, further supported by the lack of this commissure in female *T. axi* (unpublished observations, but see [11]).

Despite the minor differences, the overlapping immunoreactive patterns of several neural markers with corresponding location next to similar muscle layers of the copulatory organ support a homologous circular innervation and control of the copulatory organ. These conserved neurotransmitter patterns and putatively conserved roles in orchestration of copulation in the three closely related dinophilid species is also the foundation for more general predictions on specific function and interplay of the tested neurotransmitters (see below).

Given the strong connection of the circular innervation around the copulatory organ to the ventral nervous system in all three species, the operation of the copulatory organ is suggested to be closely relying on cerebral regulation and – input. In *D. gyrociliatus* dwarf males the results supported that the neural control is furthermore aided by direct input from posterior sensory cells (supported by [21,22]).

### Conserved roles of specific neuropeptides on muscular activity?

The similar, most likely homologous, innervation of the copulatory organ in three dinophilid species as depicted by using antibodies against specific neurotransmitters used in this study also infers a conserved functionality of these signalling molecules in regulating copulation within the family. The unpaired, medioventral glandomuscular copulatory organ most likely represents a novel formation in Dinophilidae or their close ancestors, since other described copulatory organs of meiofaunal annelids (e.g., Parergodrilidae [36,37], Syllidae [38,39], Pisionidae [40] and Hesionidae [41,42]) are mainly formed in pairs, and may be external rather than internal structures with often less muscularisation [35–46]. Although the variety of the copulatory organs and thereby possible alterations in their neural control within different annelid families suggests that the architecture of the nervous system regulating reproduction might vary, the overall functionality of the neurotransmitters on muscles, glands and other tissues may still be conserved. Neuropeptide functions on specific organ systems including the reproductive organs have not been studied in detail in other annelids but in a number of other metazoans, e.g., the model ecdysozoan animals *Caenorhabditis elegans* [28,47–49] and *Drosophila melanogaster* [50–53]. The effect of several neuropeptides on the regulation of copulatory and mating behaviour is well analysed in molluscs [54–57], which are closer related to the annelids. Neuroregulation of mating/copulation in many gastropods is however more complicated than it is expected for dinophilids, since many of the investigated mollusks are hermaphrodites, and therefore need to regulate both male and female behaviour and organ systems within one animal [54,58,59].

Many of the here tested neuropeptides, but especially FMRFamide and FVRIamide, have been detected in the penis nerve innervating the male copulatory organ in gastropods such as *Lymnaea stagnalis* and *Helix aspera* [27,54,56,60], which resembles the results of this study: we also find immunoreactivity of these neuropeptides in the ventral and dorsal penial commissures and the circumpenial fibre mass. However, the anti-FMRFamide antibody used in this study is commercially available and applicable to a broad range of animals, thereby possibly labelling several different FMRFamides. Studies in the nematode *C. elegans* have shown FMRFamide-like peptides to cause both muscle inactivation or – activation, depending on the respective peptide [28,47,61]. We therefore cannot hypothesize about the more specific functionality of these neuropeptides in dinophilids based on our data, but it is likely that the neural elements labelled by the used FMRFamide-antibody are involved in the muscular contractions of the penis sheath and possibly also of the seminal vesicles and spermioducts. FVRIamide was shown to inhibit the cycle of muscular contraction and ‒relaxation in the vas deferens of *Lymnaea-stagnalis* [27], which coincides with our findings of FVRIamide immunoreactivity in the nerves tracing the seminal vesicles, spermioducts, and penis sheath. The largely overlapping immunoreactivity patterns between FVRIamide and FMRFamide could indicate their opposing roles in coordinating the transport of sperm to the penis bulb as well as of “pumping” movements of the copulatory organ.

Myoinhibitory peptide (MIP) is less studied in molluscs, but has a rather long history of research in various insects, among them the fly *Drosophila melanogaster*, where it was found to bind to its promiscuous so called sex peptide receptor [30,62]. As such, MIP is involved in modulating behaviour according to circadian rhythms, sleep, feeding, life stage transitions, and also mating and copulation [31,52,63,64]. In contrast to our findings in dinophilids, MIP-peptides and -precursors are only present in the central nervous system (brain and ventral nerve cord), but have not been found in male accessory glands of the reproductive system or the female genital tract in *D. melanogaster* [31,52]. Remarkably, while MIP seems to act mainly myoinhibitory in insects, recent studies in larvae and juveniles of the macroscopic annelid *Platynereis dumerilii* focussing on MIP’s role in feeding and larval settlement have shown it to be myostimulatory [65–67]. The investigated species were however too young to also exhibit mating behaviour or have formed reproductive organs. The here presented study characterizes MIP-like immunoreactivity in the ventral and dorsal penial commissure, one (anterior) segmental body commissure, and the circumpenial fibre mass, but is not associated with the spermioducts or seminal vesicles, thereby separating it from the broader pattern of the previously mentioned FMRFamide- and FVRIamide-like immunoreactivity, yet indicating a more complex orchestration of the pumping movements of the copulatory organ possibly involving all of the here tested neurotransmitters. Based on the findings of especially Williams *et al.* [66], we believe that an activating role of MIP on the copulatory organ (such as demonstrated for gut musculature in larvae of a macroscopic annelid) seems to be very likely. An increase in the level of MIP in the organism would therefore also lead to an increase in the frequency of pumping movements of the penis observed in the three dinophilid species we analysed, which is similar to an increase in gut peristaltic movements in the larvae of *P. dumerilii* [66]. Future studies will prove whether the posterior sensory neurons nr. 15 in the *D. gyrociliatus* dwarf males, which are in direct connection to the circumpenial fibre mass around the penis cone, and eventually their equivalents in the larger dinophilid species, form a similar circuitry than the one described in the larvae of *P. dumerilii*. This would furthermore imply that the direct sensory-neurosecretory mechanism of MIP is not only governing life-cycle transitions (e.g. settlement and feeding [66], but can be re-deployed in other circuitries, such as the circuit underlying copulation.

Most of the neuropeptides as well as the monoamine neurotransmitter serotonin are not only found in the nervous system associated with the reproductive organs, but also in the central nervous system [11]. The majority of immunoreactive somata are located around the brain, projecting their neurites through the brain neuropil, the circumesophageal commissures, and the ventrolateral nerve cords towards the posterior end of the body [11]. It is therefore likely that the brain and the ventrolateral nerve cords are vital elements of the neural circuitry controlling mating behaviour and copulation, but which are complemented by input from local neuron accumulations such as here shown in the posterior penis ganglion and sensory neurons of *D. gyrociliatus* dwarf males.

## Conclusion

Members of the interstitial annelid family Dinophilidae use hypodermal injection for direct sperm transfer, and the glandomuscular copulatory organ developed in this family is used to both produce a hole in the epidermis of the female and to inject sperm into their bodies. This is proposedly done by the male first attaching to the female (using the adhesive glands surrounding his gonopore), then enzymatically dissolving a hole in the female’s epidermis (with secretes from the extended penis cone glands), and subsequently moving sperm first into the lumen of the penis sheath and from there pumping it into the female body [5,7,8,9]. This process requires detailed orchestration of muscular and glandular cells, which is amongst others accomplished by the nerves and ganglia in close association to the copulatory organ. Although the architecture of this innervation varies slightly between the studied males of the three dinophilid species *Trilobodrilus axi*, *Dinophilus vorticoides*, and *D. gyrociliatus*, all three species show several defined nerve bundles (commissures) embracing the copulatory organ as well as a strong connection of these structures to the ventral nervous system. The presence of immunoreactivity to all tested neurotransmitters (serotonin, FMRFamide, FVRIamide, and MIP) in one to several of these nerves strongly suggests their involvement in regulating at least the musculature of the copulatory organ and thereby also of copulatory behaviour. The conserved architectural immunohistochemical patterns across the three tested species combined with our knowledge of the animal’s behaviour (personal observations and [4,5,7,8,21,22,32]) hereby indicate a conserved functionality of these neurotransmitters across Dinophilidae. Combining our structural data with experimental data from behavioural studies in other annelids and molluscs [27,29,55], we suggest that serotonin, FMRFamide and FVRIamide act on muscles of the copulatory organ, the spermioducts, and the seminal vesicles to also support the movement of sperm from the seminal vesicles into the penis sheath [47,61,68], while especially MIP (together with the previously mentioned and other, so far untested molecules) is more strongly involved in creating peristaltic or pumping movements [27,31,65]. Although their immunoreactivity patterns by themselves cannot reveal more detailed information about their specific functionality, future studies involving the localization of the respective receptors, behavioural observations, and exposure to elevated or decreased transmitter-levels as well as comparative studies across species may unravel the intricate mechanism of neuroregulation of copulation in adult meiofaunal annelids.

## Materials and methods

### Specimens

Adult and juvenile specimens of *Trilobodrilus axi* were collected from the sandy intertidal beach of List, Sylt, Germany, from extractions of clean sand in early May and late June 2017, and kept at the Marine Biological Section, University of Copenhagen in filtered seawater (31 per mille salinity), at 15°C in the dark.

Adult specimens of *Dinophilus vorticoides* were collected from algae in the intertidal close to Kaldbak, Faroe Islands, and kept alive in 31 per mille filtered seawater at 10°C at the Marine Biological Section, University of Copenhagen for up to two months prior to fixation.

Males of *D. vorticoides* and *T. axi* were identified by means of light microscopy, anesthetized using an isotonic solution of MgCl_2_ 1:1 with filtered seawater and fixed for immunohistochemical labelling after live observations.

To obtain dwarf males of *Dinophilus gyrociliatus*, cocoons with male and female embryos still in their egg envelopes, which were supposed to hatch within the next 24h (for details about the staging, see [18]) were separated from the main culture boxes into smaller dishes in filtered seawater with 28 per mille. The cultures were originally started and maintained by Bertil Åkesson based on animals collected in Xiamen, China, and kept at the Marine Biological Section, University of Copenhagen in the dark at 20°C. Upon hatching, the dwarf males were anesthetized with isotonic MgCl_2_ 1:1 with seawater and subsequently fixed in for immunohistochemical labelling (see below).

### Light microscopy

Males of the three dinophilid species used for light microscopy were anesthetized, mounted on microscope slides and images were taken with a mounted Olympus DP73 camera mounted on an Olympus IX70 inverted compound microscope (Olympus Corporation, Tokyo, Japan, owned by K. Worsaae) in combination with the CellSens Entry software package v1.6.

### Immunohistochemistry and confocal laser scanning microscopy (CLSM)

Quadruple labelings including (immuno-)stains for/against F-actin (Alexa Fluor 488-labelled phalloidin, A12379, Invitrogen, Carlsbad, USA), DNA (405nm fluorescent DAPI, included in Vectashield), tubulin (monoclonal mouse anti-acetylated α-tubulin (T6793, Sigma, St. Louis, USA, RRID:AB_477585)), serotonin (monoclonal rabbit anti-serotonin (5-HT), S5545, Sigma, RRID:AB_477522), FMRFamide (polyclonal rabbit anti-FMRFamide, 20091, Immunostar, RRID:AB_572232), and polyclonal antibodies raised in rabbits against MIP and FVRIamide originally designed after *Platynereis dumerilii* sequences [23,65–67], and checked for their sequence similarity by Kerbl *et al.* [11]. We furthermore designed antibodies against the specific sequence of amidated *Dinophilus gyrociliatus*-MIP (CGWGGNKGMSMWamide, GenScript Biotech Corp, New Jersey, USA), and its immunoreactivity pattern is similar to the ones observed with the previously used *Platynereis*-antibody. All described and following steps were conducted at room temperature (RT) on the rocking board.

Males of the three dinophilid species were first anesthetized, followed by fixation with 3.7% paraformaldehyde in phosphate buffered saline (PBS) 1:1 with isotonic MgCl_2_-solution for 1h. This step was followed by several rinses in PBS, before the specimens were preincubated in 0.1% PTA (PBS + 0.1% Triton-X + 0.25% bovine albumin serum (BSA) + 10% sucrose + 0.05% NaN_3_) for 1h.

Following preincubation, the specimens were incubated in 0.1% PTA with primary antibodies against acetylated α-tubulin (final concentration 1:400) and either serotonin (final concentration 1:400), FMRFamide (final concentration 1:400), FVRIamide (final concentration 1:1000), or MIP (final concentration 1:1000) for 48h, and subsequently rinsed several times in PBS. This step preceded the incubation of the specimens in the secondary antibodies (goat anti-mouse antibody conjugated with CY5 (115-175-062, Jackson ImmunoResearch, West Grove, USA), and goat anti-rabbit antibody conjugated with TRITC (T5268, Sigma), final concentrations 1:800) for 24h. Subsequently, specimens were rinsed in PBS several times and incubated in 0.33μM phalloidin in 0.1% PTA for 1h. Finally, specimens were rinsed repeatedly in PBS and mounted in Vectashield (Vector Laboratories, Burlingame, USA). Specificity of primary antibody binding was tested by treating specimens only with secondary antibodies, but otherwise using the same protocol. The reproducibility of the results was checked by replicating the incubations.

The mounted specimens were examined with an Olympus IX81 inverted microscope combined with a Fluoview FV-1000 confocal unit (property of K. Worsaae at the Marine Biological Section, University of Copenhagen, Denmark). Recorded Z-stacks were imported in the Imaris 7.6 software package (Bitplane Scientific Software, Zürich, Switzerland) for further three-dimensional investigations. Snapshots were used to export individual sections and smaller stacks from Imaris for plate preparation.

### Image preparation

Brightness, saturation and contrast of LM- and CLSM-images were adjusted in Adobe Photoshop CC 2018 (ADOBE Systems Inc., San Jose, USA) prior to assembling figure plates in Adobe Illustrator CC 2018. The latter was also used to create the schematic drawings.

## Declarations

### Ethics approval and consent to participate

Not applicable

### Consent for publication

Not applicable

### Availability of data and material

All data analysed in this study are used in the tables and the figures of this article. The original image stacks can be made available after personal contact with the corresponding authors.

## Author’s contributions

AK and KW designed the project. EWT conducted immunohistochemical experiments and CLSM-analyses of the immunoreactivity patterns of *Trilobodrilus axi* and created the schematic drawings for this species. AK conducted the corresponding experiments on the two *Dinophilus*-species. AK, EWT and KW analysed the data and KW and AK wrote the manuscript. All authors read and agreed to the final version of the submitted text.

## Acknowledgments

The authors want to thank Nicolas Bekkouche and Brett Gonzalez for their help with sampling the animals used in this study, and Gáspár Jékely and Markus Conzelmann for their work on designing the antibodies against FVRIamide and MIP (designed against sequences of *Platynereis dumerilii*), providing us with aliquots to do the immunohistochemical labelling, and helping us with the production of our own antibodies.

